# When cold-bloods heat up: meta-analytical evidence that climatic variability mediates behavioural fever in amphibians and reptiles

**DOI:** 10.64898/2026.06.01.729281

**Authors:** Danilo Giacometti, Leonardo M. Servino, Laura C. Cabanzo-Olarte, Kênia C. Bícego, Carlos A. Navas

**Affiliations:** Departamento de Fisiologia, Instituto de Biociências, Universidade de São Paulo, São Paulo, SP, 05508-900, Brasil; Departamento de Ecologia, Instituto de Biociências, Universidade de São Paulo, São Paulo, SP, 05508-900, Brasil; Departamento de Morfologia e Fisiologia Animal, Faculdade de Ciências Agrárias e Veterinárias, Universidade Estadual Paulista “Júlio de Mesquita Filho”, Jaboticabal, SP, 4884-900, Brasil

**Keywords:** Amphibia, body temperature, climate change, disease, pathogens, Reptilia

## Abstract

Fever is a widespread and adaptive defence response that enhances immune performance through an increase in body temperature above normal values. In ectotherms, fever is expressed behaviourally through the selection of warmer microhabitats following infection, yet its magnitude and determinants vary widely across species and environments. Here, we performed a phylogenetically informed meta-analysis of behavioural fever in amphibians and reptiles to test whether its expression was shaped by climatic thermal variability, pathogen identity, and taxonomy. Specifically, we tested the hypotheses that (i) species from thermally variable environments would exhibit stronger behavioural fever than species from thermally stable environments, consistent with the climate variability hypothesis, and that (ii) reptiles would exhibit stronger fever responses than amphibians due to lower hydrothermal constraints. Across 47 studies encompassing 103 effect sizes, we found that behavioural fever is widespread but highly context-dependent. We found that evidence for behavioural fever was strongest in species from more thermally variable habitats, regardless of body size and phylogeny, suggesting that access to thermally heterogeneous landscapes and enhanced behavioural plasticity amplify the capacity to sustain febrile responses. Contrary to our hypothesis, amphibians exhibited stronger fever responses than reptiles, possibly reflecting differences in baseline thermoregulatory demands and environmental opportunity, or as a consequence of methodological artefacts. The expression of behavioural fever also varied with pathogen identity, with bacterial infections eliciting larger body temperature increases than fungal or viral challenges, although pathogen representation was uneven across studies. Together, our results support the idea that the capacity to express behavioural fever depends on access to thermally heterogeneous landscapes, and may vary according to pathogen biology. Ultimately, our study emphasises that temperature is not a background condition for host-pathogen interactions, but an active and environmentally contingent component of ectotherm immune defence in amphibians and reptiles.

## Introduction

Pathogens represent a pervasive selective pressure in natural systems, shaping host physiology and behaviour, population dynamics, and patterns of biodiversity (Tompkins et al. 2011). Exposure to pathogens like bacteria, fungi, and viruses is ubiquitous and unavoidable in wild populations (Gulland 1995). Consequently, hosts have evolved a diverse array of defences to mitigate the costs and consequences of infections. Host defences against pathogens range from molecular and cellular immune responses to behavioural strategies that reduce exposure, limit pathogen growth, or enhance immune efficiency (Danilova 2006; Rakus et al. 2017).

Importantly, the expression and effectiveness of these defences depend on factors like ecological context, energetic constraints, and the behavioural options available to the host (Meylan et al. 2013; Laughton et al. 2017; Beukema et al. 2021). Therefore, understanding host responses to pathogens requires an integrative framework that links immunology, physiology, behaviour, and the physical environment in which infection occurs.

Fever is a widespread and biologically important defence response. In vertebrates, fever appears as a pathophysiological reaction to contact with infectious or inflammatory agents (i.e., “exogenous pyrogens”) in which core body temperature is regulated above its typical range (IUPS 2001). The adaptive value of fever lies in its capacity to enhance immune performance and inhibit pathogen replication, making it an effective response across host-pathogen systems (Kluger 2015). Febrile responses, however, can be achieved through different means that include both autonomic (involuntary) and behavioural responses that vary among taxa (Bícego et al.2007). In mammals and birds, fever is generated endogenously through a coordinated process that involves an immune- and endocrine-induced shift in hypothalamic set-points, which is effected through thermogenic and heat-retention mechanisms such as shivering and non-shivering thermogenesis and/or peripheral vasoconstriction (Bicego et al. 2007; Gray et al. 2013; Evans et al. 2015). By contrast, amphibians and reptiles produce limited endogenous heat and generally body heat less effectively than endotherms (Tattersall et al. 2012). As a result, fever in amphibians and reptiles is expressed behaviourally through the selection of warmer microhabitats after pathogen exposure (Vaughn et al. 1974; Myhre et al. 1977). Thus, behavioural fever externalises a physiological process into behaviour, coupling immune activation, movement decisions, and habitat use (Kluger 1979; Beukema et al. 2021). Behavioural fever has been documented across several ectotherm taxa, including invertebrate and vertebrate species, and is increasingly recognised as a key component of amphibian and reptile immune defence (Myhre et al. 1977; Burns et al. 1996; Bicego et al. 2007; Sheng et al. 2025).

At the physiological level, fever emerges from a cascade of interacting processes that are generally conserved in vertebrates (Kluger 1979). Exogenous pyrogens are detected by pattern recognition receptors in immune cells, triggering the release of pro-inflammatory cytokines (e.g., Interleukin-1 ß, Tumor Necrosis Factor), which subsequently stimulate prostaglandin (e.g., Prostaglandin E2) production. These mediators alter the neural processing of thermal information and increase thermoregulatory drive, thereby inducing fever (Bernheim & Kluger 1976; Rakus et al. 2017). In amphibians and reptiles, this cascade does not result in an increase in metabolic heat production; instead, it culminates in shifts in locomotory patterns and biases habitat selection toward warmer environments (i.e., positive thermotaxis), ultimately resulting in a regulated increase of thermoregulatory set-points (*sensu* Barber & Crawford, 1977; Kluger 1977; Monagas & Gatten 1983). Thus, the expression of behavioural fever depends not only on immune signalling, but also on sensory integration, locomotor capacity, and the structure of the surrounding thermal landscape (Muchlinski 1985; Ortega et al. 1991; Boltaña et al. 2013; Ortega-Chinchilla et al. 2022). Behavioural fever should therefore be viewed as an emergent property of physiological plasticity interacting with environmental thermal conditions, rather than as a fixed trait expressed uniformly across species or contexts. Importantly, behavioural fever is typically inferred relative to temperatures selected under non-infected conditions (i.e., a reference value considered normal; hereafter “RVCN”), such that its interpretation depends on whether experimentally derived thermal preferences represent stable thermoregulatory tendencies or context-dependent responses to laboratory conditions, raising the question of how comparable behavioural fever responses are across taxa (Bícego et al. 2007; Cabanzo-Olarte et al. 2024).

Environmental thermal variability is consequently expected to play a central role in shaping behavioural fever in amphibians and reptiles. The climate variability hypothesis (CVH) posits that organisms inhabiting thermally variable environments should evolve broader thermal tolerances and greater physiological and behavioural plasticity than organisms from more thermally stable climates (Stevens 1989; Addo-Bediako et al. 2000). Comparative evidence supports this prediction across multiple ectothermic taxa, with species from more variable environments typically experiencing and tolerating greater variation in thermal biology traits (Huey & Hertz 1984; Clusella-Trullas & Chown 2014; Giacometti et al. 2024). Although the CVH has rarely been examined in the context of behavioural fever (but see Adelman et al. 2010), it predicts that species exposed to pronounced temperature fluctuations should be better equipped to exploit the thermal landscape as a regulatory tool during infection. For such species, fever-inducing temperatures may fall within the range routinely experienced during activity, potentially reducing the physiological costs of maintaining a febrile state (Muchlinski 1985; Merchant et al. 2007; Sauer et al. 2018). Conversely, species from thermally stable environments may incur greater costs when reaching or maintaining fever-inducing temperatures, particularly when environmental thermal heterogeneity is limited (Sears et al. 2016), thereby constraining the magnitude of behavioural fever they can express. Under such constraints, alternative sickness behaviours (e.g., behavioural anapyrexia, fasting, or reduced activity) may represent potential adaptive responses to infection, highlighting that the absence of behavioural fever should not necessarily be interpreted as an absence of thermoregulatory engagement (Cabanzo-Olarte et al. 2024).

Critically, the CVH should not be treated as an inherently latitudinal hypothesis, because its corollaries depend strongly on the scale at which thermal variability is quantified. Contrary to suggestions from preliminary studies, thermal heterogeneity does not increase monotonically from the tropics to higher latitudes, nor is it adequately captured by geographic coordinates alone (Aigang et al. 2009; Curtis et al. 2016). Instead, thermal variability depends on spatial (e.g., microhabitat vs. regional climate) and temporal scales (e.g., diurnal vs. seasonal shifts), as well as landscape features (e.g., elevation, canopy cover) (Sears et al. 2016; Anderson et al. 2022).

These factors are relevant for ectotherms that rely on external sources of heat to behaviourally regulate body temperature during infection. Testing the applicability of the CVH to behavioural fever therefore requires climatic metrics that capture biologically relevant thermal environments rather than coarse geographic proxies (Wu et al. 2025; Klinges et al. 2026).

Amphibians and reptiles provide an informative comparative system to test the extent to which the CVH predicts the expression of behavioural fever. These groups differ markedly in integumentary permeability, water balance, and thermoregulatory patterns (Lillywhite 2006; Angilletta 2009), all of which may influence the expression of behavioural fever. Amphibians possess highly permeable skin and generally show high evaporative water loss, thus being assumed to face strong hydrothermal trade-offs that should limit sustained warming under natural conditions, although the extent to which these constraints are expressed under experimentally permissive settings remains unclear (Tattersall 2007; Greenberg & Palen 2021; Navas et al. 2021). By contrast, reptiles generally possess less permeable integuments and maintain higher body temperatures, leading to the expectation of stronger fever responses (Lillywhite 2006; Phillips et al. 2016). Nonetheless, substantial variation exists within and across both clades (Cabanzo-Olarte et al. 2024), suggesting that environmental context and evolutionary history may be more informative predictors than taxonomy alone. Pathogen identity adds further complexity, as different pathogens vary in thermal sensitivity and immune activation pathways, potentially influencing both the magnitude and timing of fever responses (Rakus et al. 2017). Consequently, understanding behavioural fever requires integrating host physiology, pathogen biology, and thermal opportunity across ecological contexts.

Despite decades of experimental work, no quantitative synthesis has evaluated whether climatic thermal variability predicts the expression of behavioural fever across ectothermic tetrapods. At the same time, behavioural fever studies differ markedly in experimental design, pathogen exposure, and the conditions used to quantify thermal responses, raising important questions about the comparability of fever expression across taxa and ecological contexts. Here, we present a phylogenetically informed meta-analysis of behavioural fever in amphibians and reptiles to test whether fever expression is better explained by climatic thermal variability and pathogen identity than by taxonomic differences alone, and to evaluate how environmental context mediates temperature-based defence mechanisms in natural systems. We predicted that species from more thermally variable habitats would express stronger behavioural fever than those from more thermally stable habitats, and that this effect would be mediated by both taxonomy (greater effect in reptiles than amphibians) and pathogen identity after controlling for body size effects and shared ancestry. Our study gives insight into factors that may affect behavioural fever in tetrapod ectotherms, and highlights that defence mechanisms cannot be understood in isolation from the environments in which they are expressed.

## Material and methods

### Literature search and inclusion criteria

In August 2025, we conducted a systematic literature search on the *Scopus* and *Web of Science Core Collection* databases following the Preferred Reporting Items for Systematic Reviews and Meta-analyses (PRISMA) statement (Page et al. 2021). On both databases, we used search strings consisting of the following keywords: (“amphibia*” OR “reptile*” OR “frog*” OR “toad*” OR “anura” OR “salamander” OR “caudat*” OR “caecilian*” OR “gymnophiona” OR “lizard*” OR “snake*” OR “squamat*” OR “turtle*” OR “testudine*” OR “tortoise*” OR “alligator*” OR “crocodil*”) AND (“infection*” OR “disease” OR “pathogen*” OR “bacteria*” OR “fung*” OR “vir*”) AND (“body temperature” OR “Tpref” OR “Tsel” OR “Tset”) AND (“fever” OR “behavioural fever” OR “behavioral fever”) AND (“thermoregulat*” OR “therm*”). We retrieved a total of 791 documents (**Figure S1**), and after removing 127 duplicates, we assessed the eligibility of the remaining 664 studies by reading their titles and abstracts. As inclusion criteria, we considered studies that (i) used amphibians or reptiles as study systems, (ii) evaluated animals that were either experimentally or naturally exposed to an immune challenge or pathogen, (iii) stated the type of immune challenge or pathogen (e.g., bacteria or virus), (iv) stated whether animals were raised in captivity or sourced from nature, (v) compared and reported descriptive statistics on the body temperatures of healthy and infected individuals within the same study, and (vi) provided information on body mass (M_b_).

### Effect size calculation and data collation

From each study, we extracted the mean body temperatures, standard deviations, and sample sizes for both control and treated/infected groups. We obtained these data from the main text, tables, supporting information, and figures, using the *WebPlotDigitizer* software (Rohatgi 2014) for the latter. We used the standardised mean difference (Hedges’ *g*) and its associated sampling variance (V*_g_*) as a measure of effect size (Hedges 1981):

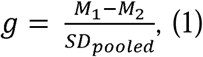

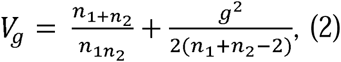

where M_1_ = body temperature of infected individuals, *M*_2_ = body temperature of control individuals, SD_pooled_= pooled standard deviation, *n*_1_ = sample size of infected individuals, and *n*_2_ = sample size of control individuals. By arranging our data as such, positive *g* values indicated that animals exhibited higher body temperatures after an infection, and negative *g* values indicated that animals sustained lower body temperatures after an infection. Thus, the greater the *g*, the greater the evidence of behavioural fever.

We recorded the following information from every study in our database: species, origin of animals (nature vs. captive), M_b_ (g), and type of infection (bacterial, fungal, denatured endogenous pyrogens, viral, or other). Importantly, we grouped studies involving nematodes, ticks, fly larvae, and heavy metals into the “other” category because these treatments did not readily fit within our predefined infection classes and were individually underrepresented in the literature. Pooling them into a broader category avoided overparameterisation and prevented the creation of moderator levels represented by a single or very small number of effect sizes.

Additionally, we recorded the latitude and longitude of where the study was conducted, although such information was not available for all studies in our data set. When necessary, we updated species names following the nomenclature available in the Amphibian Species of the World Database (Frost 2025) and the Reptile Database (Uetz et al. 2025).

### Meta-analysis, bias estimates, and sensitivity analyses

We performed all analyses using R (version 4.5.1) in RStudio (version 2025.09.2+418) and created our figures with the phytools (Revell 2012), orchaRd (Nakagawa et al. 2023), ggplot2 (Wickham 2016), and ggpubr (Kassambara & Kassambara 2020) packages. To evaluate study and experimental-level effects on the expression of behavioural fever, we fitted multilevel meta-analytical models with the “rma.mv” function from the metafor package (Viechtbauer 2010). We considered *g* as the response variable and V*_g_* as the sampling variance meta-analytical models. We included the following random terms in all models: study authorship, origin of study animals, the species’ identity (ID), the phylogenetic relatedness of the species in our data set, and an observation-level random effect (i.e., residual variance). To account for phylogenetic effects, we built a variance-covariance matrix based on recent amphibian and reptile phylogenetic trees (Zheng & Wiens 2016; Jetz & Pyron 2018). We pruned, combined, and dated these phylogenies using the ape (Paradis and Schliep 2019) and geiger (Pennell et al. 2014) packages.

To determine the overall effect size for behavioural fever across amphibians and reptiles, we fitted an intercept-only model. To understand how taxonomy and methodological attributes influenced the expression of fever, we fitted a model that considered taxonomical class (amphibians vs. reptiles), origin (nature vs. captivity), and infection type (bacterial, fungal, pyrogen, viral, or other) as moderator variables. We quantified heterogeneity (I^2^) estimates and their associated 95% confidence intervals (CIs) with the “i2_ml” function from the orchaRd package (Nakagawa et al. 2023), which allowed us to partition the total heterogeneity (I^2^_total_) across each random factor in a given model. Thus, we estimated the heterogeneity associated with the identity of the studies (I^2^_study_), the origin of animals (I^2^_origin_), the species under study (I^2^_species_), the phylogenetic relatedness (I^2^_phylogeny_), and the residual variance (I^2^_residual_). The sum of these I^2^ values amounts to I^2^_total_, which ranges from 0 to 100% (Nakagawa & Santos 2012).

Generally, 25, 50, and 75% are considered, respectively, as the thresholds for low, moderate and high heterogeneity. Low I^2^ values indicate that the observed heterogeneity is the outcome of sampling error, while high I^2^ indicate that heterogeneity is not random (Nakagawa & Santos 2012). We also estimated the phylogenetic signal attributed to our models with the phylogenetic heritability index (H^2^) (Lynch 1991). H^2^ values range from 0 to 1 and are interpreted just like Pagel’s λ (Pagel 1999). Therefore, H^2^ values close to 0 suggest that evolutionary history did not influence effect sizes, whereas values close to 1 suggest that shared ancestry has an influence on effect sizes (Nakagawa & Santos 2012).

We assessed the presence of publication bias (Rosenthal 1979) in our data set by fitting a modified version of Egger’s test (Egger et al. 1997) that had the residuals of the intercept-only meta-analytical model as the response variable and V*_g_* as the predictor. We considered an intercept that was different from zero as evidence of publication bias in the data set (Nakagawa & Santos 2012). As a complement, we used the “funnel” function from the metafor package (Viechtbauer 2010) to visually inspect the relationship between model residuals and effect size precision (i.e., 1/standard error). We also tested for the presence of time-lag bias, which occurs when the magnitude of effect sizes decreases over time as more data are collected (Jennions & Møller 2002). To do so, we fitted a linear regression that had *g* as the response variable and the year of publication of each effect size as the response variable (Nakagawa & Santos 2012).

Finally, we performed sensitivity analyses to evaluate model fit and robustness. First, we assessed model fit through a visual inspection of the distribution of residuals. Second, we estimated Cook’s distance (*D*) with the “cooks.distance.rma.mv” function from the metafor package (Viechtbauer 2010), which computes how removing a certain data point influences a given model. We considered *D* values greater than 1 as extreme (Viechtbauer 2020). Third, we computed DFBETAS with the “dfbetas.rma.mv” function from the metafor package (Viechtbauer 2010) to evaluate how the presence or absence of an influential data point influenced model parameters. We considered DFBETAS values greater than 1 as influential cases (Viechtbauer & Cheung 2010). When necessary, we refitted models and compared model outputs to determine whether the removal of possibly influential effect sizes changed the qualitative interpretation of our results.

### Bioclimatic data and phylogenetic analyses

Since the relationship between thermal variability and geographical coordinates depends strongly on spatial and temporal scales, and extends beyond the simplistic view that higher latitudes experience greater thermal variation than lower latitudes (Curtis et al. 2016; Anderson et al. 2022), we avoided treating latitude as a proxy for thermal variability. Instead, we extracted site-specific climatic variables associated with thermal variability from Climatologies at High Resolution for the Earth’s Land Areas (CHELSA) (Karger et al. 2017) at a spatial resolution of 30 arsec (∼1 km). To maximise biological relevance, we restricted these analyses to studies that sourced animals from nature and provided the geographical coordinates for their collection sites. We used the sp (Pebesma & Bivand 2005) and raster (Hijmans et al. 2013) packages to obtain coordinate-specific data on mean diurnal range (BIO2), isothermality (BIO3), temperature seasonality (BIO4), annual temperature range (BIO7). The mean diurnal range is computed as the average difference between monthly maximum and minimum air temperatures, isothermality quantifies the relative contribution of day–night variation to annual thermal variation, temperature seasonality is given by the standard deviation of mean monthly temperatures, and the annual temperature range is a measure of the amplitude between the warmest and coldest months (Karger et al. 2017; Brun et al. 2022).

We then assessed correlations among climatic variables to minimise collinearity and identify biologically meaningful predictors of behavioural fever. Mean diurnal range and isothermality emerged as suitable candidates because other thermal variables were strongly correlated with one another (**Figure S2**). For mean diurnal range, higher values pertain to sampling localities characterised by greater day–night temperature fluctuations, whereas isothermality describes the extent to which daily thermal variation contributes to overall annual climatic variability. Values close to 1 pertain to sampling localities with consistent daily temperatures relative to the annual thermal cycle, while values close to 0 indicate larger annual temperature variation compared to the daily range. Thus, these variables capture complementary dimensions of thermal heterogeneity potentially relevant to thermoregulatory decision-making during infection.

To evaluate whether species originating from thermally variable habitats exhibited greater evidence for behavioural fever, we fitted Bayesian phylogenetic generalised linear mixed models (PGLMMs) with the “pglmm” function from the phyr and INLA packages (Rue et al. 2017; Li et al. 2020). For the global model, we considered the difference between infected and control body temperatures as the response variable, and mean diurnal range (scaled and mean-centred), isothermality (scaled and mean-centred), and M_b_ (log_10_-transformed) as the predictor variables. Because multiple effect sizes were available for some species and sampling effort was uneven across taxa, we included species identity as a random effect and incorporated phylogenetic covariance to account for non-independence among observations and shared evolutionary history. We used a complexity penalising prior to automatically determine the best prior distribution for our global model (Simpson et al. 2017) and we also computed Pagel’s λ (Pagel 1999) as a measure of phylogenetic signal.

To determine whether climatic variables improved explanatory power, we compared the global model with alternative models containing different combinations of predictors (seven models total). We performed model selection based on the deviance information criterion (DIC), in which higher DIC weight (DIC*_w_*) values indicate better model fit (Spiegelhalter et al. 2002). We assessed the robustness of our best-ranked model through a null-modeling approach that compared whether our observed R^2^ value exceeded values expected by chance (Hawkins et al. 2017). We generated 1,000 null data sets in which our response variable was randomly permuted while retaining the original predictor variables and phylogenetic structure. We refitted each permutated data set using PGLMMs, and extracted R^2^ values to create a null distribution. We compared the empirical R^2^ to the null R^2^ distribution to assess whether the explanatory power of our best-ranked model exceeded or was similar to that obtained by chance.

## Results

### Data set

In total, 47 studies passed our inclusion criteria and we were able to calculate 103 effect sizes for 49 species of amphibians and reptiles (**Figure 1**). Most of the data we obtained came from reptiles (66 effect sizes for reptiles vs. 37 effect sizes for amphibians), with Squamata being the most studied order (89.39% of effect sizes), followed by Testudines (7.58% of effect sizes) and Crocodilia (3.03% of effect sizes); we did not obtain any data from Sphenodontia. In amphibians, Anura was the most studied order (91.89% of effect sizes), followed by Caudata (8.11% of effect sizes); we did not retrieve any data from Gymnophiona. Species origin was balanced between captive-bred (53 effect sizes) and naturally-sourced (50 effect sizes) individuals. We obtained exact geographical coordinates for all studies conducted on species sourced from nature. Our final data set spans six decades of research on behavioural fever (from 1970s to 2020s), with most studies being published in the 2010s (**Figure S3**).

**Figure 1.**
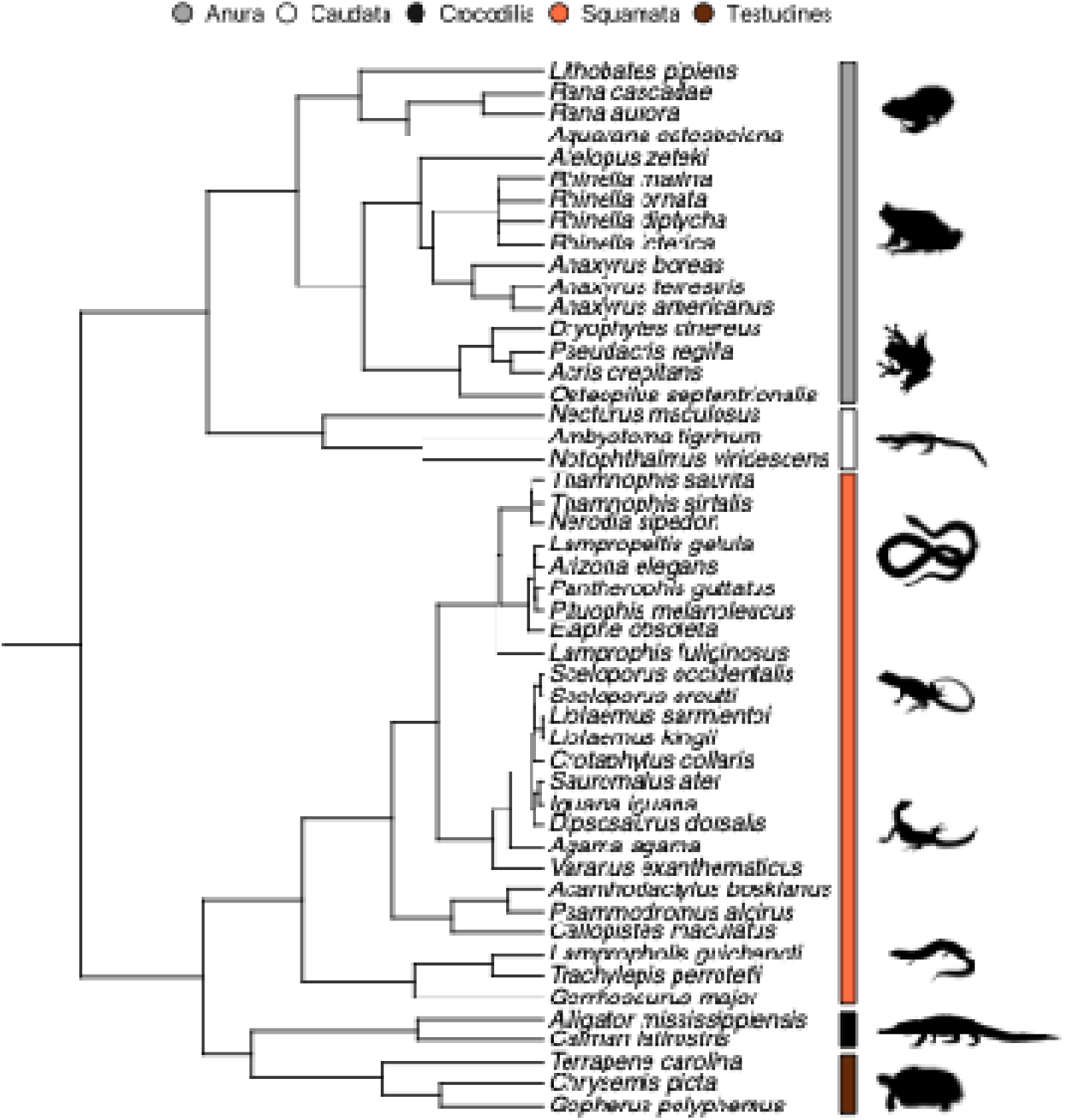
Phylogenetic relationship among the species of amphibians and reptiles represented in our data set. This tree was created based on two time-calibrated phylogenies of amphibians (Jetz & Pyron 2018) and reptiles (Zheng & Wiens 2016). Amphibian and reptile orders are colour-coded, with Anura in grey, Caudata in white, Crocodilia in black, Squamata in orange, and Testudines in brown. We obtained silhouettes that represent major amphibian and reptile taxa from PhyloPic’s (https://www.phylopic.org/) public repository.

### Behavioural fever differs between amphibians and reptiles, and depends on infection type

We present our meta-analytical results as effect size estimates ± standard error (SE) [95% confidence intervals (CIs)]. Based on the intercept-only model, we found that amphibians and reptiles, on average, exhibited behavioural fever when infected (*g* = 0.94 ± 0.42 [0.10, 1.79], *p* = 0.01) (**Figure 2**). We also found that amphibians showed greater evidence of behavioural fever than reptiles (amphibians: *g* = 2.38 ± 1.07 [0.25, 4.51], *p* = 0.02; reptiles: *g* = 0.81 ± 1.01 [−1.17, 2.80], *p* = 0.41) (**Figure 3a**), with no contribution of whether animals were sourced from nature or captivity (*g* = −0.13 ± 0.79 [−1.72, 1.45], *p* = 0.86). Moreover, animals infected with bacteria (*g* = 1.45 ± 0.69 [0.08, 2.82], *p* = 0.04) and infecting agents in the “other” category (*g* = 1.63 ± 0.81 [0.03, 3.23], *p* = 0.04) were more prone to exhibit behavioural fever than those infected with virus (*g* = 0.73 ± 0.95 [−1.15, 2.61], *p* = 0.44), fungi (*g* = 0.26 ± 0.75 [−1.23, 1.75], *p* = 0.73), or denatured endogenous pyrogens (*g* = 1.30 ± 0.88 [−0.45, 3.05], *p* = 0.14) (**Figure 3b**). I^2^ values from both models were high, suggesting that most of the observed variation remained unexplained by random terms (**Table 1**). There was no evidence of phylogenetic signal in neither meta-analytical model (**Table 1**).

**Figure 2.**
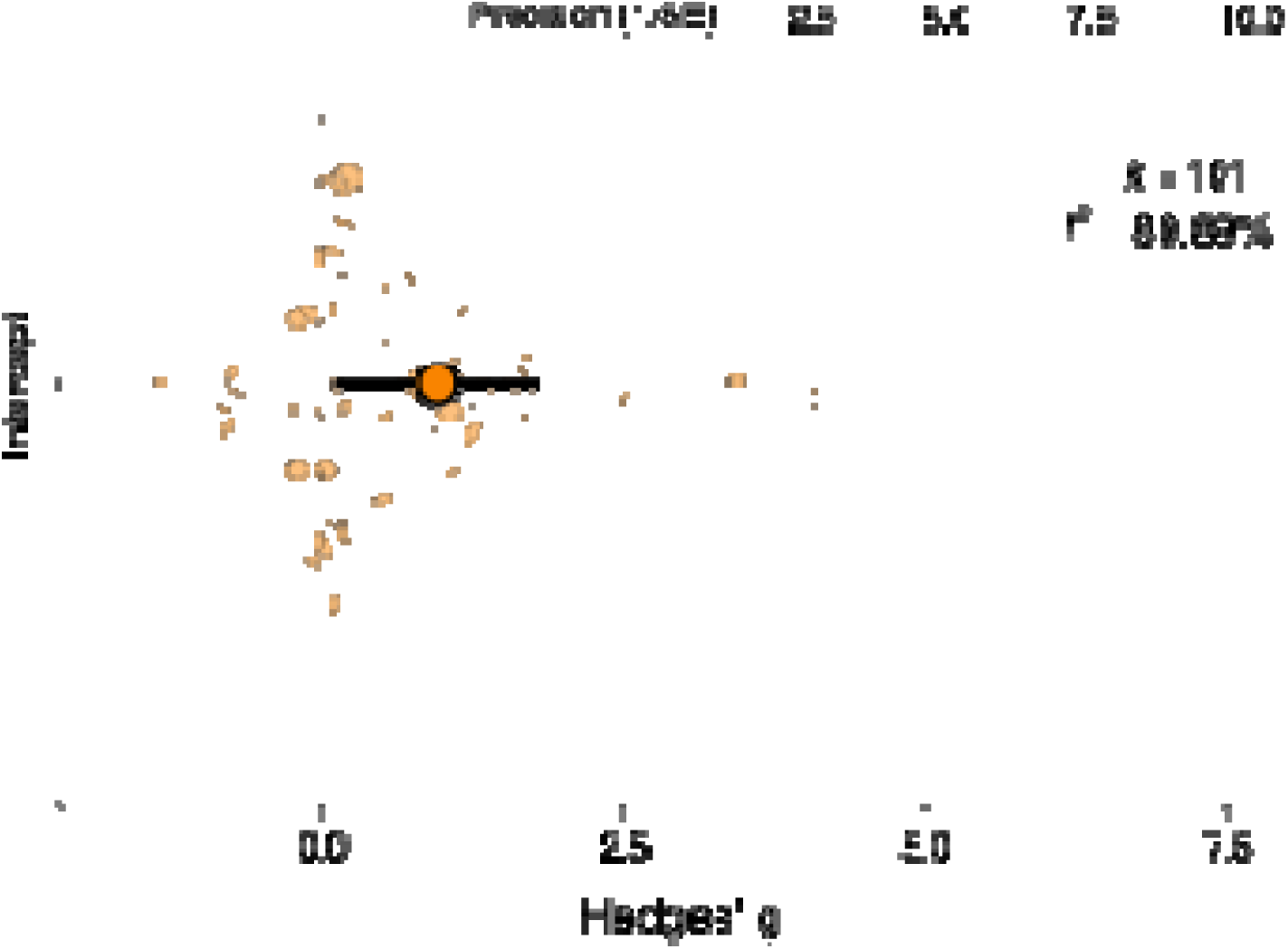
The intercept-only meta-analytical model supports the presence of an overall behavioural fever response in amphibians and reptiles, as evidenced by a positive *g* estimate and 95% confidence intervals (CIs) that do not overlap the zero. The large orange dot represents the model estimate, and the horizontal line depicts the 95% CI. Semi-transparent dots represent individual effect sizes, with dot size representing effect size precision. The vertical dotted lines denote the zero. *k* = sample sizes. *I^2^*= total heterogeneity.

**Figure 3.**
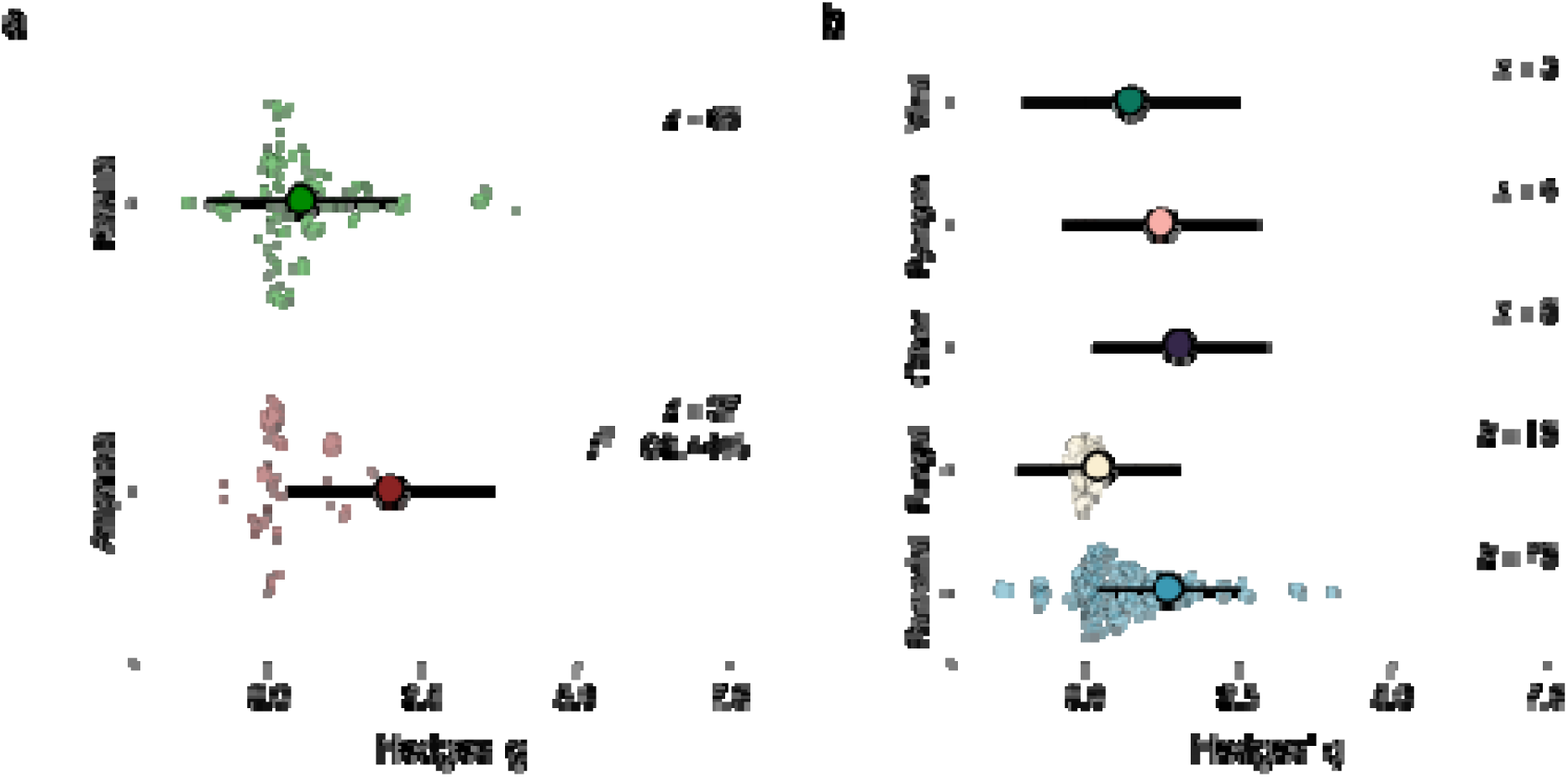
**a**. Amphibians display stronger evidence of behavioural fever than reptiles. **b**. Infections by bacteria and other agents (nematodes, ticks, fly larvae, and heavy metals) are more prone to cause behavioural fever responses than infections by fungi, endogenous pyrogens, or viruses. In both panels, large dots represent parameter estimates, and the horizontal lines depict the 95% confidence intervals. Semi-transparent dots show individual effect sizes, with dot size representing effect size precision. The vertical dotted lines denote the zero. *k* = sample sizes. *I^2^* = total heterogeneity.

**Table 1.** Heterogeneity (I^2^) and phylogenetic heritability index (H^2^) estimates for multi-level meta-analytical models. Values are reported as estimates with 95% confidence intervals (CIs) in parentheses.

| Random term | Intercept-only model | Taxonomy and infection type model |
| --- | --- | --- |
| Study | 6.62 (2.52, 47.20) | 1.19 (0, 35.85) |
| Origin | 0.01 (0, 16.89) | 10.62 (10.13, 13.41) |
| Species | 30.86 (1.32, 27.06) | 19.20 (11.08, 29.72) |
| Phylogeny | 30.62 (1.32, 27.06) | 44.60 (11.08, 49.72) |
| Residual | 22.51 (16.95, 46.99) | 17.61 (9.17, 37.96) |
| Total | 90.63 (85.35, 92.70) | 93.24 (85.59, 94.39) |
| $H^2$ | 0.45 (0, 1.92) | 0.25 (0, 1.92) |

We found evidence of both publication bias (β = −2.28 ± 0.34 [−2.97, −1.60], *p* < 0.001) (**Figure S4**) and time-lag bias (β = 53.74 ± 19.38 [15.27, 92.20], *p* = 0.006) (**Figure S5**) in our data set. Our sensitivity analyses indicated the presence of three possibly influential data points in our meta-analytical models (i.e., DFBETAS > 1). Accordingly, we removed these data from our data set and refitted our models. Doing so did not alter the qualitative interpretation of our results (see Supplementary Code).

### Daily thermal variation predicts behavioural fever

We report PGLMM results by providing posterior estimates and their corresponding 95% CIs. Species that experienced a wider diurnal thermal range displayed greater shifts in body temperature when infected (posterior estimate = 1.13 [0.37, 1.89], R^2^ = 0.68). Because diurnal thermal range was mean-centred and scaled, the posterior estimate indicates that an increase of 1 standard deviation in daily thermal variability (approximately 2.50°C in our data set) was associated with a 1.13°C difference in body temperature. In raw units, this translates to an approximate 0.45°C increase in body temperature for every 1°C increase in mean diurnal range. This effect was not mediated by isothermality (posterior estimate = −0.05 [−1.24, 1.15]) nor M_b_ (posterior estimate = 0.41 [−0.68, 1.48]) and was associated with a relatively low phylogenetic signal (λ = 0.32) (**Figure 4**). Random terms from this model suggested a modest effect for phylogeny (variance = 1.42 [0.38, 11.73]) and residual variation (variance = 3.32 [1.97, 6.30]), with no evidence of residual autocorrelation (Ljung-Box Q Test; Q = 6.08, *p* = 0.73). Our observed R^2^ value did not overlap with the randomly generated R^2^ distribution (**Figure S6**), supporting the robustness of our model.

**Figure 4.**
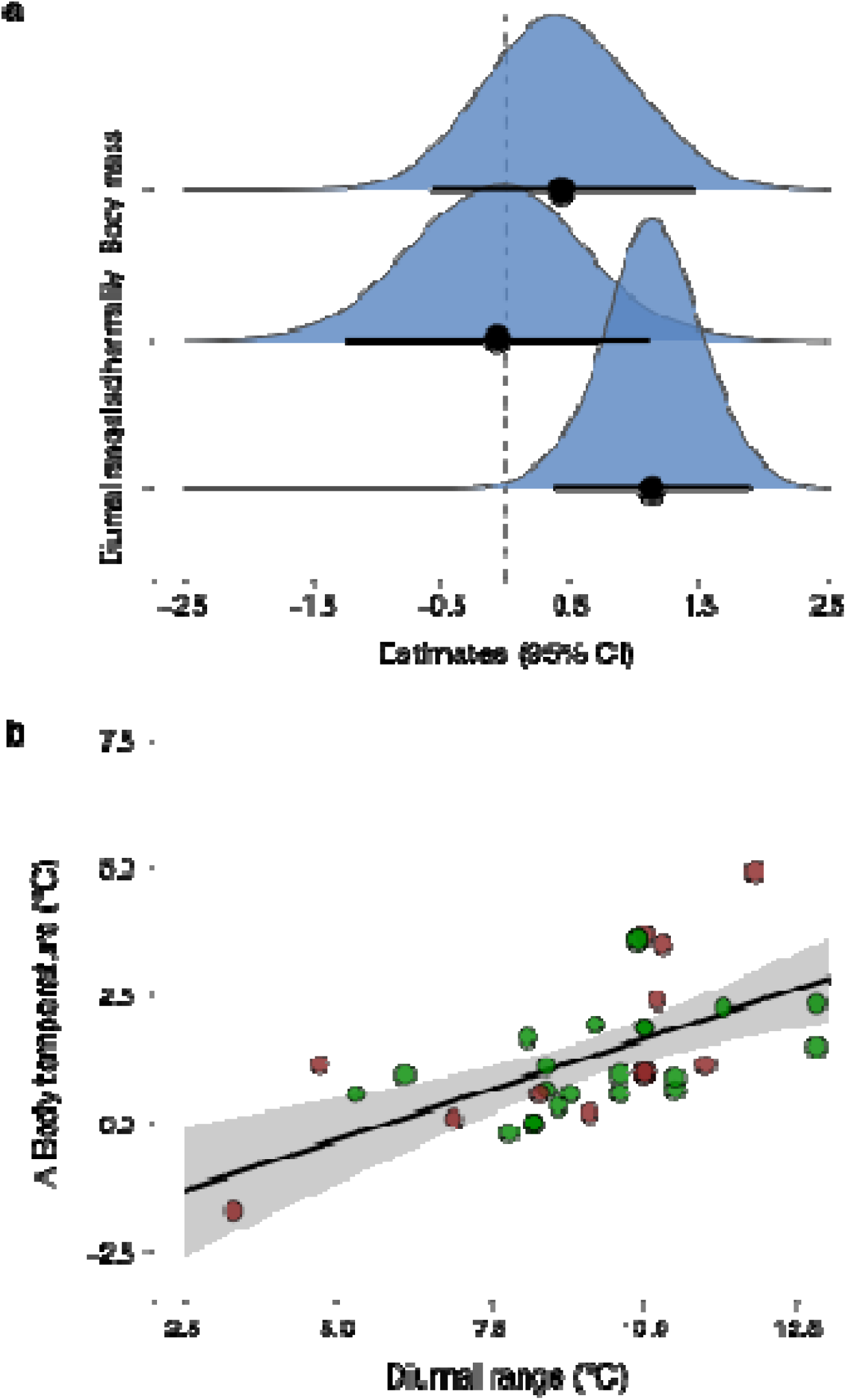
**a**. Posterior distributions for fixed terms from the Bayesian phylogenetic generalised linear mixed model evaluating whether species from thermally variable habitats showed greater evidence of behavioural fever. Diurnal range was the only significant model parameter. Black dots represent model estimates and the black horizontal bars show the 95% confidence intervals (CIs). Densities for the posterior distributions are represented in light blue for each model parameter. The vertical dashed line marks the zero. **b**. Relationship between shifts in body temperature (Δ body temperature) following an infection and mean diurnal range. Species from habitats that are more thermally variable on a daily basis exhibit greater evidence for behavioural fever than those from more stable habitats (R^2^ = 0.68). Dots are colour-coded (red = reptiles, green = amphibians) and represent individual data points. The black horizontal line shows the predicted relationship between Δ body temperature and mean diurnal range, and the grey shaded area represents the 95% CI.

## Discussion

### Behavioural fever is widespread but context dependent

Our meta-analysis supports the idea that infection is frequently associated with positive thermoregulatory responses across amphibians and reptiles, consistent with behavioural fever as traditionally defined in the literature. This finding is consistent with work demonstrating that amphibians and reptiles can behaviourally elevate body temperature following an immune challenge (reviewed in Kluger 1979; Bicego et al. 2007; Cabanzo-Olarte et al. 2024). The positive overall effect size across amphibians and reptiles reinforces the view that diverse behavioural responses to temperature constitute a central component of amphibian and reptile biology, including from an immunological standpoint. However, the high heterogeneity observed across models indicates that thermal responses are not uniform. Instead, it suggests that additional aspects such as ecological and experimental context must be considered to properly understand the factors that shape fever expression in amphibians and reptiles. Indeed, rather than representing a fixed trait, behavioural fever appears to result from the interaction between conserved immune signalling pathways, the thermal opportunities afforded by the surrounding landscape, and the experimental contexts through which thermoregulatory responses are quantified (Kluger 1979; Beukema et al. 2021; Ortega-Chinchilla et al. 2022). In other words, our data suggest that although immune activation may bias thermoregulatory drive toward warmer environmental temperatures via cytokine and prostaglandin signalling pathways (Bícego et al. 2007), the realisation of a febrile response, in species that do display this response, should depend largely on the capacity to behaviourally exploit the thermal landscape (Merchant et al. 2007). This interpretation aligns with conceptual frameworks that emphasise the role of habitat heterogeneity and climatic variability in shaping thermoregulatory responses in tetrapod ectotherms (Sears et al. 2016; Logan et al. 2019).

### Why do amphibians show stronger behavioural fever than reptiles?

Contrary to our prediction, we found that amphibians exhibited stronger evidence of behavioural fever than reptiles. Although amphibians are often assumed to face stronger hydrothermal constraints than reptiles because of their highly permeable skin and elevated evaporative water loss (Wells 2007), our results suggest that such constraints may be context dependent rather than absolute. In amphibians, behavioural thermoregulation experiments are typically conducted under hydrated and thermally permissive laboratory conditions, often relying on humid thermal gradients that incur low hydrothermal costs (Navas et al. 2021). Under such conditions, the hydric costs typically associated with sustained warming in nature may be reduced, potentially allowing amphibians to express stronger evidence of heat-seeking behaviours. In turn, this draws attention to a longstanding issue in comparative physiology: what animals can do under experimentally permissive conditions and what they routinely do in nature (Waddle et al. 2024; Laterza-Barbosa et al. 2025). Accordingly, the stronger behavioural fever observed in amphibians may reflect physiological capacity expressed under permissive laboratory conditions rather than the realised thermoregulatory behaviour observed in the field (but see Waddle et al. 2024). Future work would benefit from explicitly comparing laboratory and field body temperatures during infection, while also manipulating hydration state and thermal opportunity to determine when hydrothermal trade-offs constrain the expression of behavioural fever in amphibians.

Differences in baseline thermoregulatory demands between amphibians and reptiles may further contribute to our observed pattern. Indeed, many species of reptiles maintain higher body temperatures and function within narrower temperature ranges than amphibians, particularly among active thermoregulators (Angilletta 2009). Consequently, infected reptiles may already operate near temperatures that optimise immune performance (Bradshaw 1988), which may reduce the scope for additional warming to be detected as behavioural fever. In such cases, fever may be expressed through subtle adjustments in basking, activity timing, or microhabitat use rather than through large increases in mean body temperature. Although these thermoregulatory responses are better documented in reptiles than in amphibians (Angilletta 2009; Navas et al. 2021), the use of pre-post comparisons of body temperature likely does not capture such fine-scale adjustments. Alternatively, amphibians and reptiles can also exhibit regulated decreases in body temperature (i.e., anapyrexia) under certain contexts (e.g., hypoxia), which have been documented as distinct thermoregulatory states alongside fever (Bícego et al. 2007). While empirical evidence for behavioural anapyrexia following infection is still limited (Deen & Hutchison 2001), evidence from other contexts suggests that maintaining cooler body temperatures might be an adaptive strategy to conserve energy and mitigate physiological stressors (Tattersall & Boutilier 1997; Bícego et al. 2007; Herczeg et al. 2025). In this sense, the absence of an overt fever response in some reptiles may reflect a shift toward cooler body temperatures rather than a lack of positive thermotaxis, underscoring the need for future work to explicitly test whether infection can elicit alternative temperature adjustments and behaviours (Cabanzo-Olarte et al. 2024).

A third, non-mutually exclusive explanation for the stronger behavioural fever observed in amphibians compared to reptiles involves methodological differences among studies in terms of how infections are induced, how thermal responses are quantified, and the conditions under which such responses are evaluated. Infection protocols vary widely in the literature, encompassing different classes of pathogens (e.g., bacteria, fungi, viruses, parasites), doses, routes of exposure (e.g., injection, immersion, oral inoculation, or natural exposure), and timing of temperature measurements relative to infection (Cabanzo-Olarte et al. 2024). Such variation is likely to influence both the magnitude and temporal dynamics of immune activation and, consequently, whether behavioural fever or other sickness behaviours are expressed (Cabanzo-Olarte et al. 2024). In addition, experimental approaches used to quantify thermal preference or selection differ substantially (see Navas et al. 2021), ranging from linear or circular thermal gradients of varying breadth and resolution to semi-natural enclosures with varying degrees of thermal heterogeneity (Noronha-de-Souza et al. 2015; Figueroa-Huitrón et al. 2019; Giacometti & Tattersall 2024). Although thermal gradient design, spatial complexity, and thermal resolution influence the range of temperatures available to animals (Reiser et al. 2013; Navas et al. 2021; Laterza-Barbosa et al. 2025), experimental duration may also shape the detectability of fever responses, as studies range from short-term assays lasting only a few hours (e.g., Parris et al. 2004) to multi-day experiments capable of capturing sustained thermoregulatory adjustments (e.g., Ortega et al. 1991). Additional heterogeneity arises from the techniques used to measure body temperature (e.g., cloacal thermometry, implanted data loggers, infrared thermography), which differ in temporal resolution, invasiveness, and physiological relevance (Taylor et al. 2021). A deeper issue, however, concerns whether laboratory-based measurements of behavioural thermoregulation carry the same biological meaning across amphibians and reptiles.

Indeed, behavioural fever, as traditionally defined, is quantified relative to a baseline thermal state (i.e., RVCN) (see Cabanzo-Olarte et al. 2024). A central question is, thus, whether the RVCN reflects a strong thermoregulatory preference that is repeatable across experimental settings, or whether it is largely contingent on the species under study and the conditions imposed by each experiment (Navas et al. 2021). In heliothermic lizards, for example, temperatures selected in laboratory thermal gradients often approximate those maintained in the field (Angilletta 2009), such that experimental control temperatures may reflect meaningful thermal set-points. Under these circumstances, the RVCN tends to be robust, repeatable, and consistently thermotactic (Goulet et al. 2017). In many amphibians, however, a comparable RVCN may not exist in the same sense as a stable thermal state, as body temperature can vary continuously according to hydration state, microhabitat conditions, activity level, and local thermal opportunity (Brattstrom 1963, 1979), or even according to how animals explore the experimental arena (Navas et al. 2021; Laterza-Barbosa et al. 2025). Moreover, thermotaxis is not commonly tested in amphibians, meaning that temperatures recorded under laboratory conditions may sometimes reflect exploratory behaviour or avoidance of thermal extremes rather than a stable thermal preference (Navas et al. 2021; Giacometti & Tattersall 2024).

Consequently, whether behavioural fever can be meaningfully quantified depends not only on the magnitude of thermal change after exposure to a pathogen, but also on whether the reference thermal state reflects a pervasive thermoregulatory tendency or a variable thermal behaviour contingent on the experimental context. This distinction suggests that equivalent thermal shifts may not carry the same biological meaning across amphibians and reptiles, and that the stronger behavioural fever observed in amphibians could partly reflect greater context dependence in the thermal reference state used to quantify fever. Future work should move beyond simply documenting thermal shifts following infection and explicitly evaluate the biological meaning of the thermal reference states used to infer behavioural fever. This will require validating RVCNs through repeated measures and/or control experiments (Navas et al. 2021; Laterza-Barbosa et al. 2025), field estimates of body temperature, and independent tests of thermotactic consistency before laboratory-derived thermal shifts are interpreted as evidence of fever (Giacometti & Tattersall 2024). More broadly, advancing this field will require greater standardisation of experimental designs, clearer reporting of infection protocols and thermal environments, and stronger integration between laboratory and field studies to understand when temperature functions as a meaningful line of defence against infection.

### Pathogenic and climatic context affect behavioural fever

Our data suggest that the observed variation in the expression of behavioural fever across amphibians and reptiles reflects the influence of both pathogen identity and climatic variability. Pathogens can differ markedly in their thermal sensitivities and the immune pathways they activate, which can influence the magnitude and timing of febrile responses (Shapiro & Cowen 2012; Rakus et al. 2017; Roncarati et al. 2025). In our study, bacterial infections were more likely to elicit behavioural fever than fungal or viral infections, consistent with evidence that bacterial challenges can induce strong pro-inflammatory responses mediated by exogenous pyrogens that promote febrile states (Kluger 2015). In contrast, certain fungi may tolerate or even benefit from elevated temperatures (Muller 1956), while viral infections may cause more variable thermal responses (Woese 1960). This pattern is consistent with the thermal tolerance mismatch hypothesis, which posits that the adaptive value of sustaining elevated body temperatures depends on the relative thermal performance curves of host and pathogen (Carvalho et al. 2024; Greenspan et al. 2017). In this scenario, febrile responses should be favoured when they cause a disproportional impairment on pathogen performance relative to host function.

However, fever may be absent when pathogens tolerate or benefit from elevated temperatures. It is important to note, however, that bacterial effect sizes were disproportionately represented in our data set (*k*_bacteria_ = 75, *k*_fungus_ = 15, *k*_other_ = 6, *k*_pyrogen_ = 4, *k*_virus_ = 3), possibly because of the seminal papers that set the stage for the study of behavioural fever in tetrapod ectotherms (e.g., Vaughn et al. 1974; Bernheim & Kluger 1976). As such, although our findings suggest that behavioural fever is not a generic response to infection, it is imperative that future studies test potential fever responses across a broader diversity of pathogens and infection contexts before general conclusions about pathogen-specific thermal strategies can be drawn.

Beyond pathogen identity, our results also demonstrate that climatic thermal variability predicts the expression of behavioural fever after accounting for body size and phylogenetic effects. Species from habitats with greater day–night thermal variation exhibited larger shifts in body temperature following infection, supporting the CVH and comparative work showing that amphibians and non-avian reptiles from thermally heterogenous habitats exhibit broader thermal tolerances and enhanced behavioural flexibility (Huey & Hertz 1984; Addo-Bediako et al. 2000; Clusella-Trullas & Chown 2014). This suggests that even modest increases in day–night temperature variation can amplify the capacity of amphibians and reptiles to exhibit behavioural fever during infection, potentially enhancing immune performance in species inhabiting thermally variable environments relative to those from thermally stable habitats. We further highlight that this relationship was detected using high-resolution climatic data rather than relying on geographic proxies, reinforcing the idea that biologically relevant measures of thermal variability are essential for properly understanding patterns of thermoregulatory plasticity (Anderson et al. 2022; Giacometti et al. 2024). In this context, the scale at which thermal variability is quantified is central (Klinges et al. 2026). From an organismal perspective, diurnal thermal variation directly constrains short-term behavioural decisions and has immediate relevance for fever expression (Huey & Pianka 1977; Peterson 1987), whereas broader seasonal metrics may be less informative for acute immune responses. The relatively weak phylogenetic signal detected in our models supports the idea that the ability to realise behavioural fever is contingent on the prevailing environmental conditions, despite the fact that fever pathways are deeply conserved across the animal tree of life (Kluger 1979; Bícego et al. 2007).

### Biases and limitations

As with any broad-scale study, our conclusions must be interpreted in light of potential sources of bias. We detected evidence of publication bias and time-lag bias (Rosenthal 1979; Jennions & Møller 2002), which is consistent with broader patterns in ecological and evolutionary quantitative syntheses (Nakagawa & Santos 2012; Nakagawa et al. 2022).

However, our sensitivity analyses supported the robustness of our main conclusions, as evidenced by the fact that removing potentially influential effect sizes did not alter the qualitative interpretation of our results. Nevertheless, we reiterate that behavioural fever is inherently difficult to quantify across disparate experimental designs, and effect sizes may conflate physiological capacity with behavioural opportunity. Furthermore, our analyses focused on increases in body temperature following infection but did not explicitly account for alternative sickness behaviours (*sensu* Cabanzo-Olarte et al. 2024), like behavioural anapyrexia, which may be a biologically meaningful defence strategy in certain contexts (Bícego et al. 2007). In this vein, future research should benefit from experimental designs that formally integrate pathogen identity, thermal landscape structure, and temporal dynamics of thermoregulatory responses, thereby allowing fever and alternative sickness behaviours to be evaluated as complementary components of temperature-mediated immune defence.

### Implications in a changing climate and closing remarks

Together, our findings highlight that temperature-mediated defence mechanisms cannot be understood independently of the environments in which they are expressed. The ability to express behavioural fever depends on access to thermally heterogeneous landscapes that allow amphibians and reptiles to elevate body temperature when infected (Waddle et al. 2024), and this reliance makes such defences inherently sensitive to environmental change. Climate change is altering not only mean temperatures but also the magnitude, predictability, and spatial structure of thermal variability, often leading to thermal homogenisation at local scales (Diffenbaugh & Field 2013; Sears et al. 2016). In habitats wherein fine-scale thermal heterogeneity is reduced (e.g., through land-use change, canopy loss), the behavioural capacity to exploit temperature as an immune resource may be hampered even if physiological fever pathways remain intact. Indeed, experimental evidence suggests that infected lizards may die when infected but precluded from behaviourally raising their body temperatures (Bernheim et al. 1978). In a broader context, species that historically relied on behavioural fever may experience increased susceptibility to pathogens if their habitats undergo thermal homogenisation in future climatic conditions.

From a conservation perspective, our findings underscore the importance of preserving thermal heterogeneity as a component of disease resilience in natural populations. Microhabitats that provide warm refugia, such as sun-exposed areas, heterogeneous substrates, or structurally complex vegetation, may help sustain febrile states and improve host outcomes during infection (Waddle et al. 2024). Conversely, management actions that simplify thermal landscapes may inadvertently constrain an important line of defence against pathogens. Therefore, incorporating thermal ecology into conservation planning could enhance population persistence by maintaining the environmental conditions that allow amphibians and reptiles to behaviourally regulate body temperature during infections, especially in areas facing both climate change and emerging infectious diseases (Sauer et al. 2018; Cocciardi & Ohmer 2024). Thus, our results lend support to the view that predicting disease vulnerability under climate change will require explicit consideration of how altered thermal landscapes interact with host physiology and behaviour, and pathogen biology (Beukema et al. 2021). Ultimately, our study demonstrates that temperature is not merely a background condition for host-pathogen interactions, but an active and environmentally contingent component of amphibian and reptile immune defence.

## Funding

DG was funded by Fundação de Amparo à Pesquisa do Estado de São Paulo (FAPESP) Postdoctoral Fellowship (#2025/06560-5). LMS was funded by a FAPESP Doctoral Fellowship (#2021/10039-8). LCCO was funded by a FAPESP Postdoctoral Fellowship (#2023/12446-5). KCB was funded by a FAPESP Thematic Grant (#2021/10910-0), CAN was funded by a FAPESP Thematic Grant (#2024/13478-0).

## Author contribution

DG and CAN conceived the study. DG and LCCO collected the data. DG performed the analyses with input from LMS. DG led the writing of the manuscript. All authors contributed critically to the final version of the manuscript.

## Conflict of interest

None.

## Data availability statement

Our data and code can be accessed from: https://doi.org/10.5281/zenodo.20490561.

## Notes

https://doi.org/10.5281/zenodo.20490561

### Competing Interest Statement

The authors have declared no competing interest.

### Summary of Updates

Made some pre-submission tweaks to the introduction and discussion.

