## Supplementary material for "When cold-bloods heat up: meta-analytical evidence that climatic variability mediates behavioural fever in amphibians and reptiles" (from another version of this preprint: version 1, 2026-06-04)

**Supplementary figures 1–6**

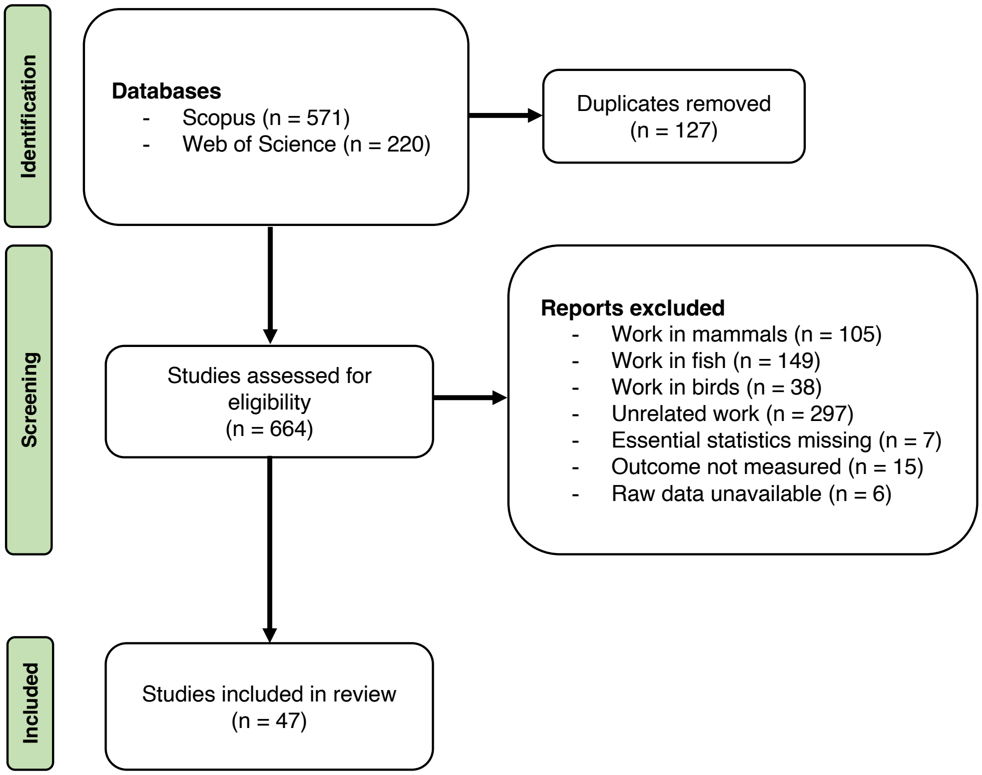


**Figure S1.** Preferred Reporting Items for Systematic Reviews and Meta-analyses Statement (PRISMA) diagram showcasing the decisions made during the building of our data set.

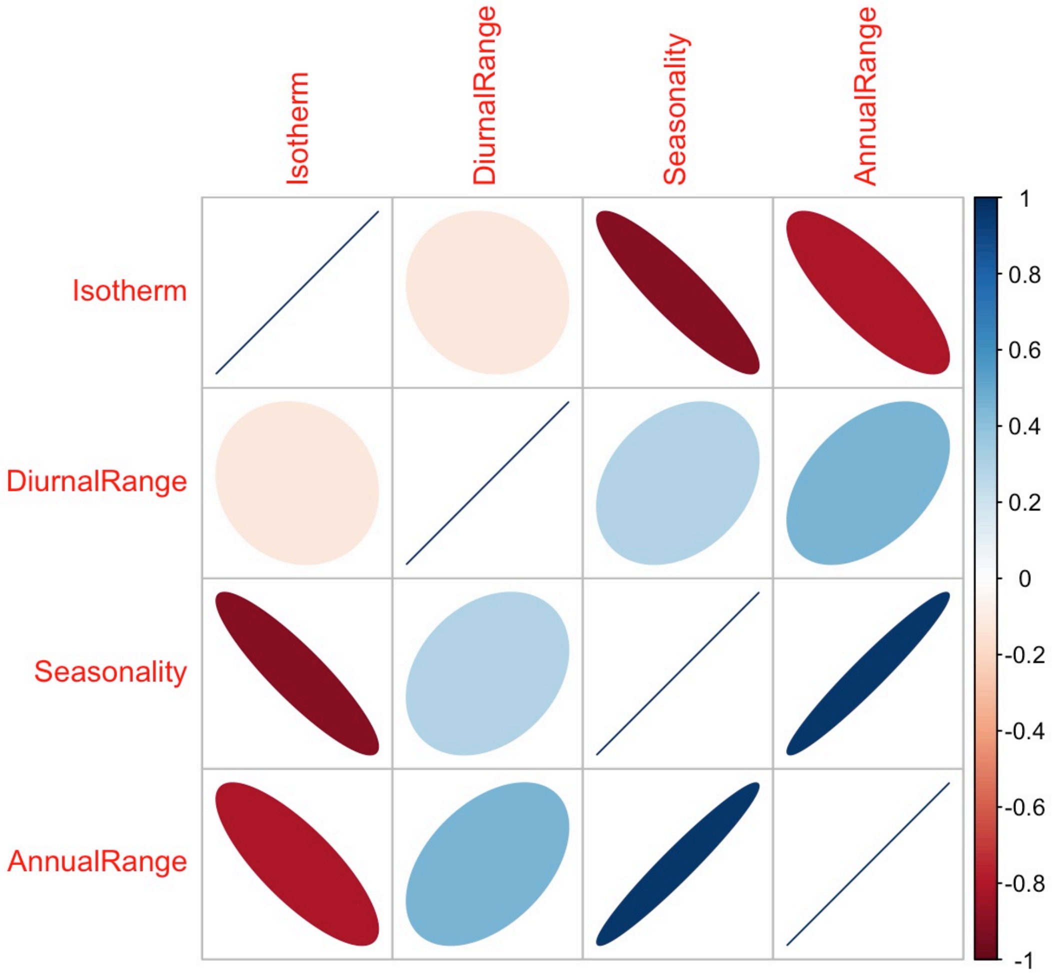


**Figure S2.** Correlation matrix of variables associated with habitat thermal variability. Size and colour intensity are proportional to the value and magnitude of the Pearson correlation coefficient, with blue indicating positive correlations and red indicating negative correlations. Based on this matrix, we selected mean diurnal range and isothermality as predictor variables in our model, since all other variables were either positively or negatively correlated.

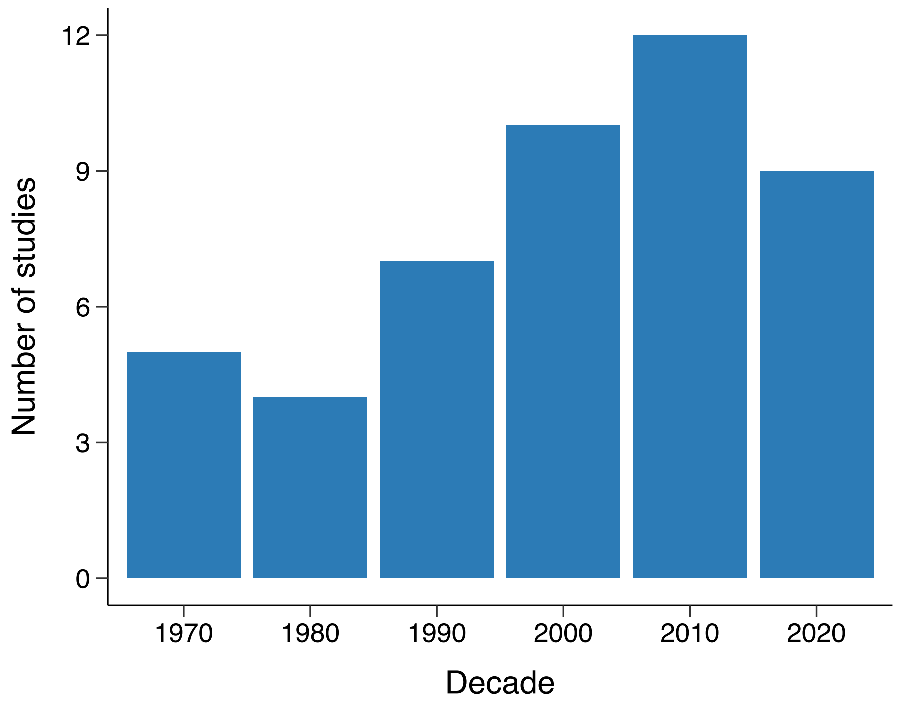


**Figure S3.** Number of studies about behavioural fever in amphibians and reptiles published per decade. Bars represent the total number of unique studies (i.e., author-year combination) included in our data set.

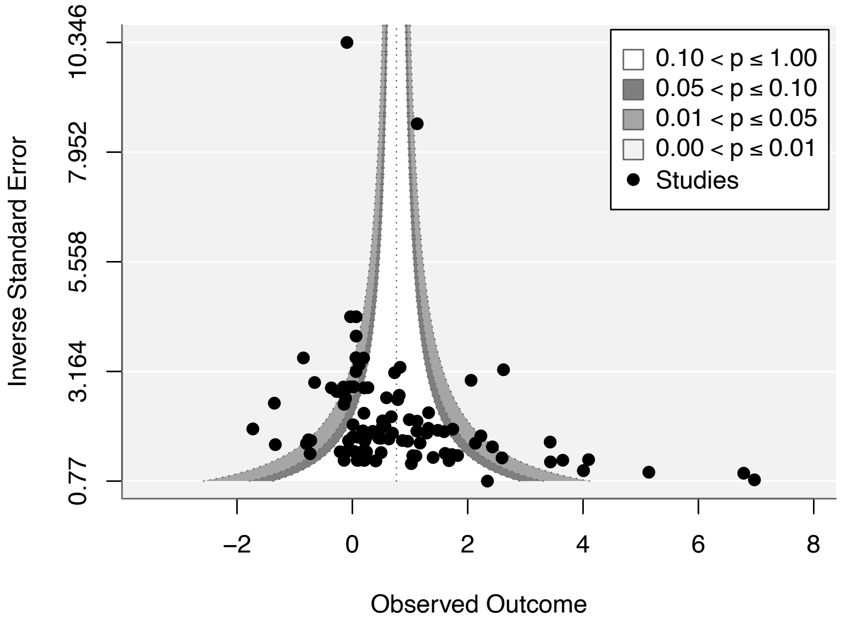


**Figure S4.** Funnel plot for the intercept-only meta-analytical model. Data are presented as model residuals as a function of the inverse of the standard errors. Shaded areas are colour-coded and represent pseudo confidence intervals (white = 90%, dark grey = 95%, and light grey = 99%).

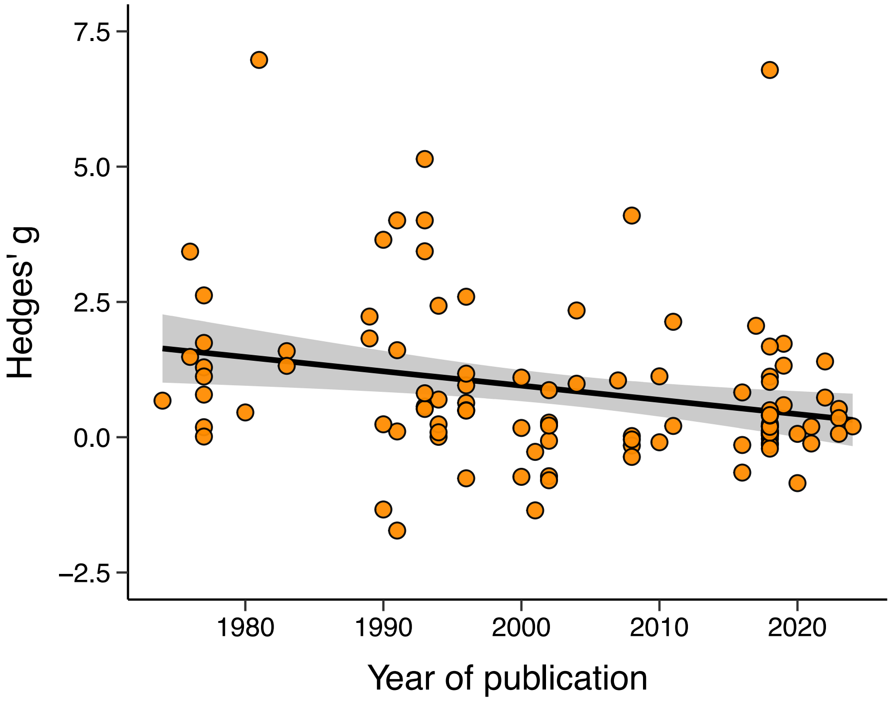


**Figure S5.** Time-lag bias plot. Publication year influenced effect sizes, with smaller effect sizes obtained over time (β = 53.74 ± 19.38 [15.27, 92.20], $p =$ 0.006).

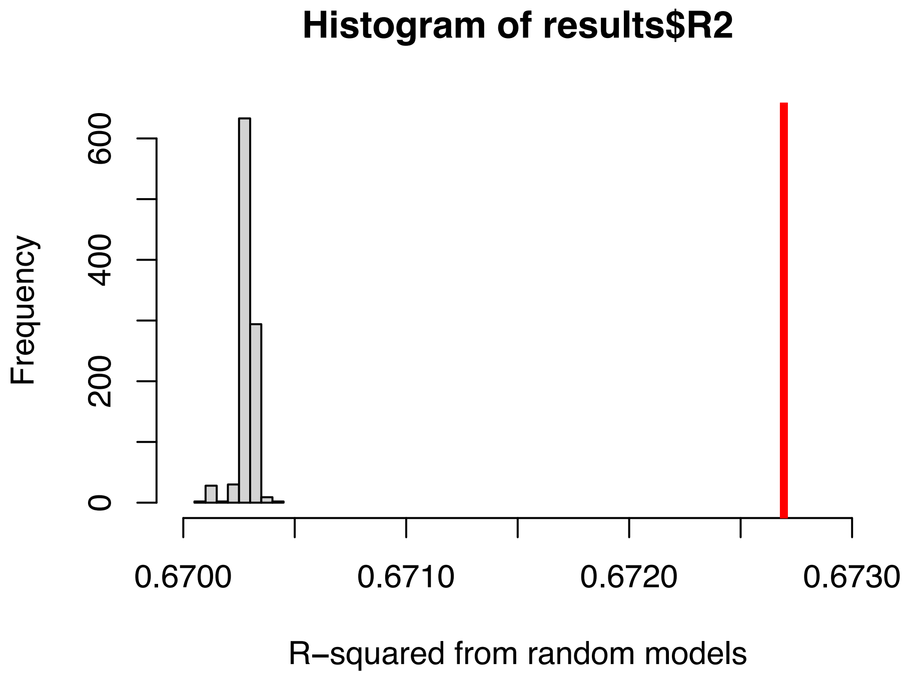


**Figure S6.** Comparison of $R^2$ values obtained from 1,000 randomly generated Bayesian phylogenetic generalised linear mixed models (grey bars) relative to the $R^2$ value obtained from our best-ranked model (red bar). Since the observed $R^2$ value did not fall within the random distribution, the explanatory power of our model is greater than that obtained by chance.
